# Diffusible Signal Factor signaling regulates multiple functions in the opportunistic pathogen *Stenotrophomonas maltophilia*

**DOI:** 10.1101/343590

**Authors:** Shi-qi An, Ji-liang Tang

## Abstract

*Stenotrophomonas maltophilia* is a Gram-negative bacterium commonly isolated from nosocomial infections. Analysis of the genome of the clinical *Stenotrophomonas maltophilia* isolate K279a indicates that it encodes a diffusible signal factor (DSF)-dependent cell-cell signaling mechanism that is highly similar to the system previously described in phytopathogens from the genera *Xanthomonas* and *Xylella*. Here we demonstrate that in *S. maltophilia* strain K279a, DSF signaling regulates factors contributing to virulence, biofilm formation and motility of this important opportunistic pathogen.

## Introduction

*Stenotrophomonas maltophilia* is a Gram-negative bacterium that is found ubiquitously in the environment and which has become an important opportunistic pathogen (Ryan et al., 2009; Berg and Martinez, 2015; An and Berg, 2018). *S. maltophilia* infections occur in cystic fibrosis and burns patients and are common in individuals with compromised immune systems who are inclined to opportunistic infections (Ryan et al., 2009; Berg and Martinez, 2015; An and Berg, 2018). The organism is commonly isolated from clinical specimens and is involved in urinary and respiratory tract infections, endocarditis and in catheter-related bacteraemia and septicaemia (Ryan et al., 2009; Berg and Martinez, 2015 An and Berg, 2018). Isolates are resistant to the majority of clinically useful antibiotics making the treatment of *S. maltophilia* infections problematic (Crossman et al., 2008). A number of laboratories have been addressing the molecular bases for virulence and antibiotic resistance in *S. maltophilia* (Ryan et al., 2009; Alvai et al., 2013; Huedo et al., 2015; Berg and Martinez, 2015; An and Berg, 2018).

*S. maltophilia* is related to bacteria from the genera *Xanthomonas* and *Xylella* (Minkwitz and Berg, 2001). In these plant pathogens, cell-cell signaling mediated by molecules of the diffusible signal factor (DSF) family control virulence factor synthesis and virulence to plants as well as the interaction of *Xylella* with its insect vector and the development of biofilms and adhesion in both genera (reviewed in Ryan et al., 2015). DSF family signal molecules are *cis*-unsaturated fatty acids, the first of which to be identified was *cis*-11-methyl-2-dodecenoic acid from *Xanthomonas campestris* (Wang et al., 2004). The synthesis and perception of the signals require products of the *rpf* gene cluster; DSF synthesis is dependent on RpfF, which has some amino acid sequence similarity to enoyl-CoA hydratases, whereas DSF perception involves a two-component regulatory system, including the complex sensor RpfC and response regulator RpfG (Barber et al., 1997; Wang et al., 2004; An et al., 2013; Ryan et al., 2015). In *Xanthomonas campestris*, the *rpfG* and *rpfC* genes are co-transcribed as the *rpfGHC* operon, although RpfH has no clear role in signaling and is not conserved in other *Xanthomonas* species or in *Xylella*. The *rpfB* gene, which encodes a long-chain fatty acyl coenzyme A ligase, is linked to *rpfF* in *X. campestris* and has been implicated in DSF turnover (Zhou et al., 2015). The relatedness of *S. maltophilia* to these plant pathogens prompted us to examine this organism for the existence and role of a DSF-dependent signaling system. In 2007, it was reported that mutation of *rpfF* had effects on different phenotypes in *S. maltophilia* (Fouhy et al., 2007). However, this paper was recently retracted due to errors in data presentation (Fouhy et al., 2018). Here we report on the outcomes of repeated key experiments that indicate the pleiotropic nature of *rpfF* mutation and show that DSF signaling controls factors contributing to virulence, motility and biofilm formation of this nosocomial pathogen.

## Materials and Methods

### Bacterial Strains and Growth Conditions

Strains and plasmids used during this study are described in Table 1. For the majority of experiments, NYGB medium was used as growth media for *S. maltophilia* strains, which contains 20 g/L glycerol (Sigma-Aldrich, UK), 3 g/L yeast extract (Difco, UK) and 5 g/L bacteriological peptone (Oxoid, UK). The measurement bacterial biofilm formation was carried out in L medium, which comprises of sodium chloride, 5 g/L; yeast extract, 5 g/L; Bactotryptone (Difco, UK), 10 g/L and D-glucose (Sigma-Aldrich, UK), 1 g/L.

**Table 1.**
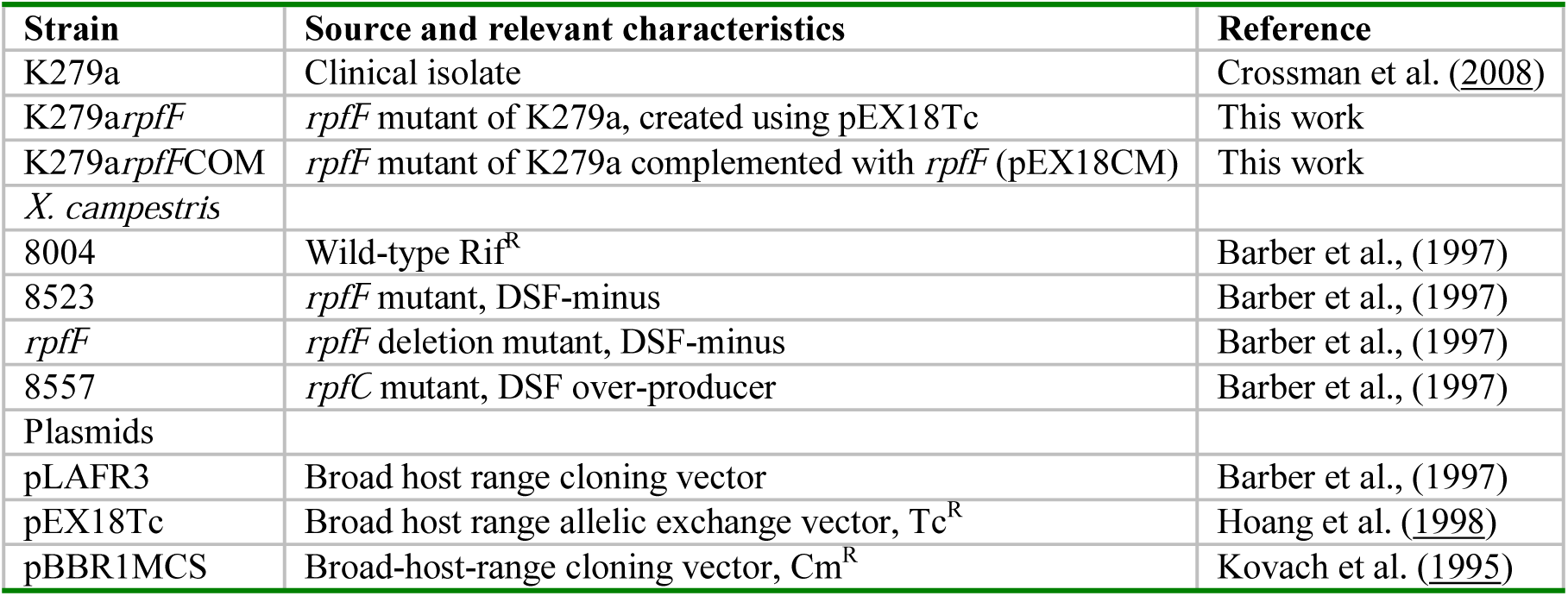
Bacterial strains and plasmid used in this study

The strain of *S. maltophilia* used was K279a (Crossman et al., 2008). In order to create a disruption of the *rpfF* gene in *S. maltophilia*, an internal fragment of the gene was amplified using the primers PEX18RPFF-F:

5’-TGACATCGTCGACGACTACCAGC-3’) and PEX1 8RPFF-R: 5’-GGCTTTCCTTGATCACCTGT-3’ and was cloned into the TOPO (Invitrogen) vector. This fragment was excised with *Eco*RI and ligated into the suicide plasmid pEX18Tc. This construct was introduced into *S. maltophilia* K279a by triparental mating. The mating mixture was plated on NYGA medium containing tetracycline (125 µg ml –1) to select for mutants. Candidate strains were analyzed by colony PCR using the primers Con-F: 5’-TTGCGTATTGGGCGCTCTTCC-3’ and Con-R: 5’-ACGATGATCGGCCTGTCGCT-3’ to confirm disruption of the *rpfF* gene by the suicide vector. For complementation studies, the *rpfF* gene was cloned into pBBR1MCS (Kovach et al., 1995).

For the complementation of *X. campestris* strains, the *rpfF* gene from *S. maltophilia* K279a with its promoter was amplified by PCR using the primers RPFFCOMF: 5’-GAATTCAGACGGCGGGGTCTTT-3’ and RPFFCOMR: 5’-AAGCTTTCAGGCCGGGTCGCCATT-3’ and the DNA fragment cloned into the TOPO vector. The *rpfF* gene was excised as a *Eco*RI –*Hin*dIII fragment and ligated into pLAFR3 cut with the same enzymes. This resulting construct was introduced into selected *X. campestris* strains by triparental mating.

### DSF extraction and synthetic DSF

DSF was extracted into ethyl acetate from culture supernatants of strains grown in NYGB as described by Barber and colleagues (1997). DSF was assayed by measuring the restoration of endoglucanase activity to an *X. campestris rpfF* mutant strain 8523 (Table 1) by extracts from culture supernatants (Barber et al., 1997). Synthetic DSF from *X. campestris* (*cis*-11-Methyl-2-dodecenoic acid) was purchased from Merck (Sigma-Aldrich, UK).

### Extracellular enzymes assays

For measurement of protease activity, strains were grown in NYG medium, 3 µl of the overnight cultures (OD600 nm≈1.0) were spotted onto an NYG plate containing 1% (w/v) skimmed milk and allowed to dry before growth at 30°C for 24(h). For measurement of endoglucanase activity, strains were grown in NYG medium overnight. Enzyme activity in cell-free culture supernatants were measured by radial diffusion assays into substrate-containing agar plates using carboxymethyl cellulose (CMC) as substrate (Barber et al., 1997).

### Motility assays

Bacterial motility assays were carried out on NYGB media that was solidified using 0.6% Eiken agar (Eiken Chemical, Tokyo). A sterile 200-µl tip was used to inoculate *S. maltophilia* strains to the centre of the plate. Plates were visualized after incubated at 30°C for 48 h.

### Biofilm formation assay

Bacterial strains were assessed for biofilm formation by aggregation in L medium as described previously (Dow et al., 2003). Here log-phase-grown bacteria were diluted to OD600 nm = 0.02 in L media broth, and 5 ml was incubated at 30°C for 24 h in 14 ml glass tubes.

### Virulence assay

Virulence was tested in *Galleria mellonella* larvae (An and Tang, 2018), which were stored at 4°C in wood shavings. *G. mellonella* were injected with 10 μl of successively diluted bacteria (1 × 10^6^ CFU). Infected *G. mellonella* were placed on Whatman paper lined Petri dishes and incubated at 37°C. The *G. mellonella* were monitored for their survival after a 24 h period. Four separate tests were conducted consisting of 10 larvae for each strain. The control groups for each experiment consisted of *G. mellonella* injected with PBS alone.

### Results and Discussion

Initial evidence for the occurrence of the DSF signaling system in *S. maltophilia* clinical isolate K279a was provided by bioinformatic analysis. Interrogation of the genome sequence of this organism (http://www.sanger.ac.uk/Projects/S_maltophilia/) using with the RpfF amino acid sequence of *X. campestris* in tBLASTn revealed a homolog of RpfF. Further analysis of a DNA sequence of approximately 8Kb (to include flanking genes) indicated the presence of an *rpfBFCG* gene cluster, related to that found in *X. campestris* (Fig. 1). No homologue of *rpfH* was identified in *S. maltophilia* (Fig. 1). The *S. maltophilia* proteins showed very high amino acid sequence similarity to their homologues in *X. campestris*; in BLASTP comparisons, E values were lower than 10^-127^.

**Fig. 1.**
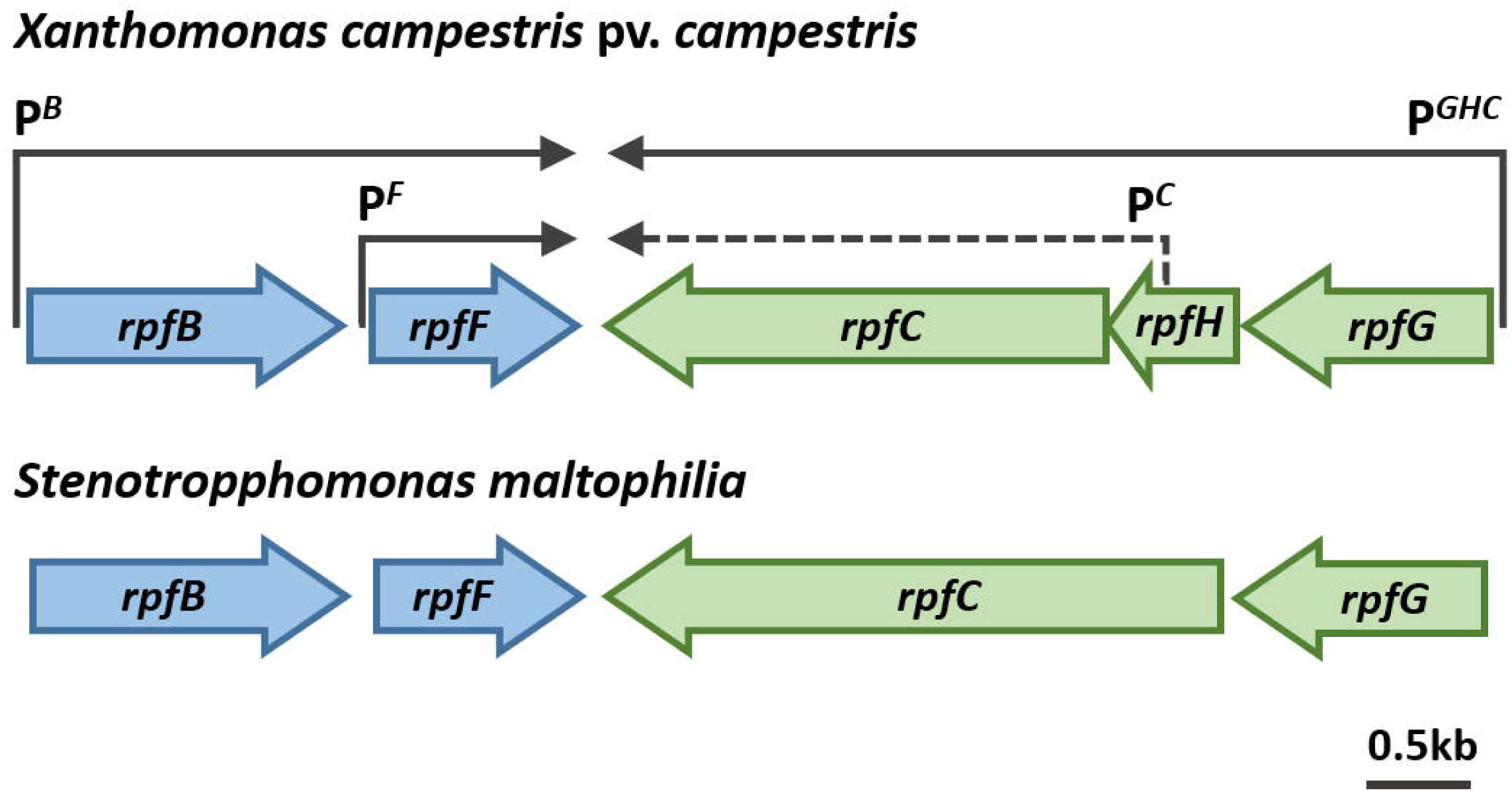
Physical map of the part of the *rpf* gene cluster from *rpfB* to *rpfG* in *Xanthomonas campestris* and *Stenotrophomonas maltophilia* K279a. The organization of ORFs predicted by sequence analysis together with predicted directions of transcription are indicated by the broad arrows. The positions of the experimentally determined transcriptional start sites in *X. campestris* together with the predicted transcripts are shown as single arrows. The sequence data for *S. maltophilia* were produced by the *S. maltophilia* K279a Sequencing Group at the Sanger Institute and can be obtained from http://www.sanger.ac.uk/Projects/S_maltophilia/).

These bioinformatic studies were supported by experimental studies to examine the production of DSF by the K279a strain. DSF in extracts of culture supernatants was assayed by measuring the restoration of endoglucanase activity to an *X. campestris rpfF* mutant (see Materials and Methods). Using this bioassay, DSF activity was detected in culture supernatants of *S. maltophilia* K279a (Fig. 2). Further evidence that the product of the *S. maltophilia* K279a *rpfF* gene directs DSF production was obtained from experiments in which the cloned gene was introduced into the *rpfF* mutant of *X. campestris*. The transconjugant produced detectable DSF production and the production of the extracellular enzymes endoglucanase and protease was concomitantly restored (Fig. 2). Furthermore, inactivation of *rpfF* in *S. maltophilia* K279a by use of the pEX18Tc suicide vector (see Materials and Methods) led to a loss of DSF synthesis as (Fig. 2).

**Fig. 2.**
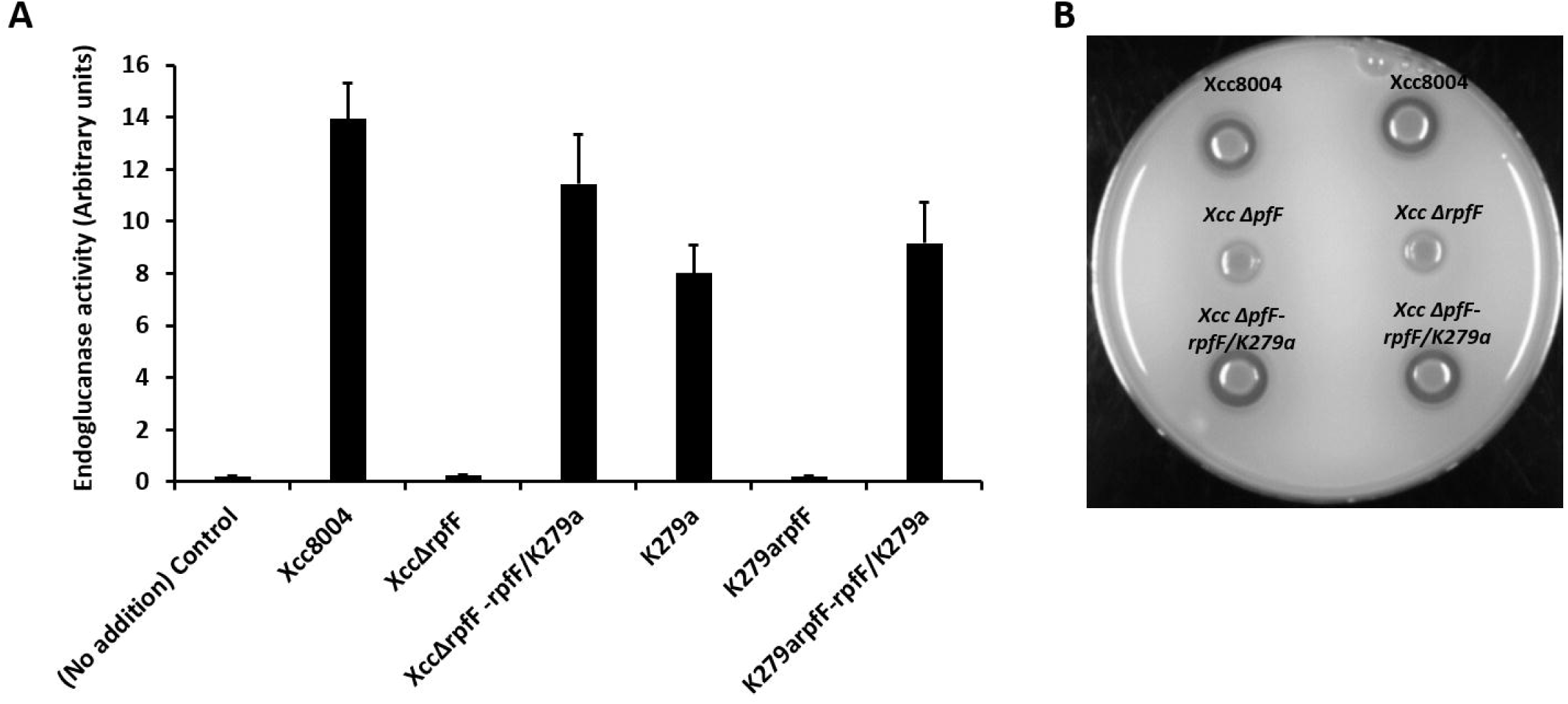
A: DSF activity in culture supernatants of strains of *S. maltophilia* K279a and *X. campestris*. Extracts were assayed using a *Xanthomonas* bioassay in which restoration of endoglucanase activity to an *rpfF* mutant is measured (A). Introduction of the *rpfF* gene from *S. maltophilia* K279a restores the synthesis of protease to the *rpfF* mutant of *X. campestris* (B). Enzyme activity was assessed by zones of clearing produced after growth of bacteria on skimmed milk agar plates. Top panel: Protease production in *X. campestris* wild type (*Xcc* 8004). Middle panel: Protease production in *X. campestris rpfF* mutant (*Xcc* Δ*pfF*). Right panel: Protease production by *Xcc* Δ*pfF* carrying the *S*. *maltophilia rpfF* gene cloned in pLAFR3.

The effects of disruption of DSF signaling through inactivation of *rpfF* in *S. maltophilia* K279a were pleiotropic. The *rpfF* mutant had severely reduced motility (Fig. 3), decreased levels of extracellular protease (Fig 3) and formed aggregates or biofilms when grown in L medium (Fig. 3). Importantly, *in trans* expression of the *rpfF* gene in the mutant could restore phenotypes towards wild type in all cases.

**Fig 3.**
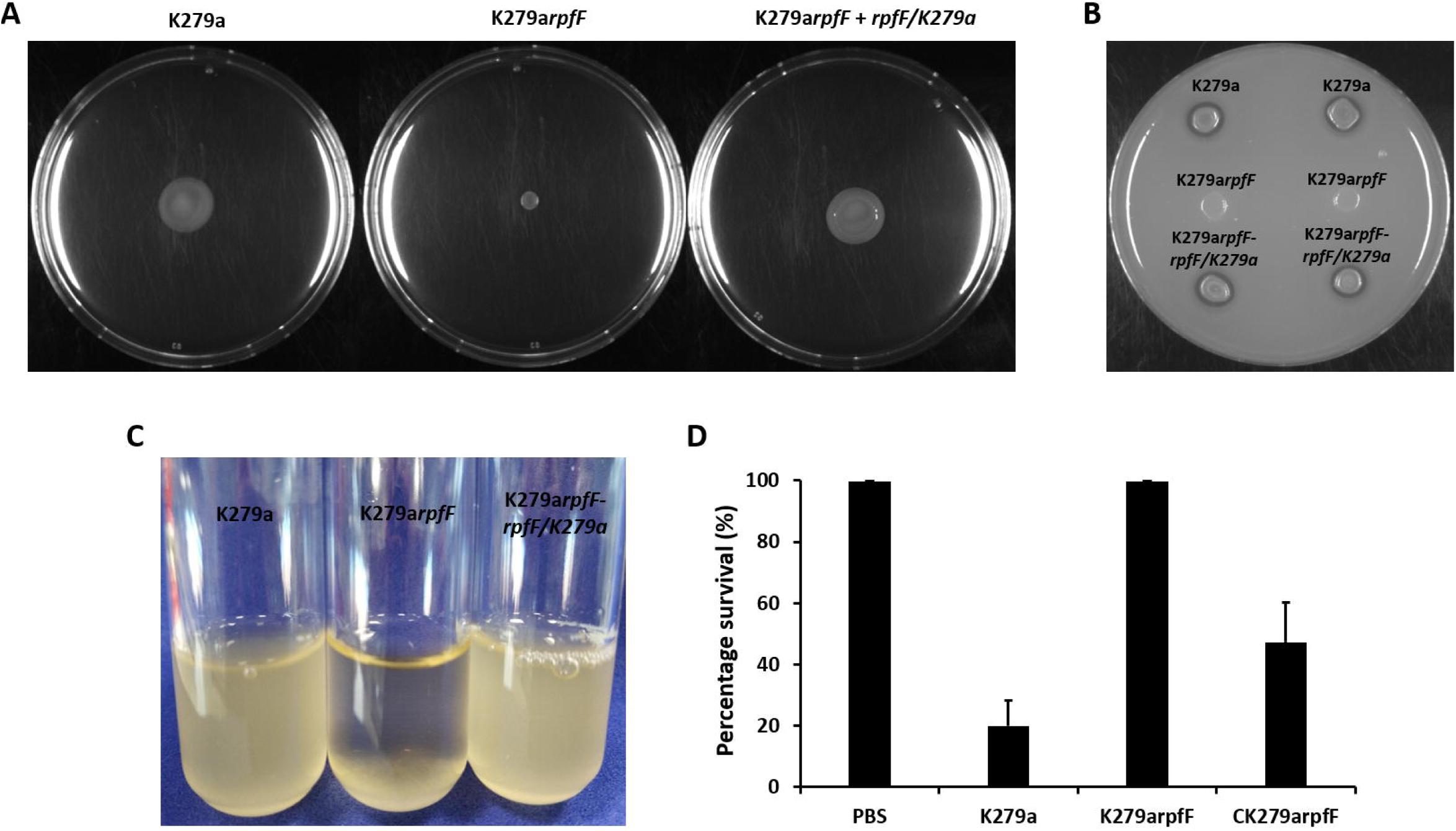
Loss of DSF signaling through mutation of *rpfF* has a pleiotropic effect in *S. maltophilia*. The *rpfF* mutant shows reduced swimming motility in 0.6% Eiken agar (A); reduced production of extracellular protease (B); aggregation when grown in L medium (C), where the wild-type grows in a dispersed fashion; (D) ability to cause disease to *Galleria mellonella* (Wax moth larvae). *In trans* expression *rpfF* gene in the *S. maltophilia rpfF* mutant could restore phenotypes towards wild-type in all cases.

The above findings demonstrated the impact of DSF signaling on aggregative behavior and protease synthesis in *S. maltophilia*, which are functions that are believed to be involved in virulence. This motivated us to test the effect of *rpfF* mutation on *S. maltophilia* virulence using the *Galleria mellonella* larvae model (see Materials and Methods). Under the assay conditions used, the wild-type *S. maltophilia* K279a killed all of the larvae after 24h, whereas the *rpfF* mutant of *S. maltophilia* K279a produced no killing (Fig. 3). These findings suggest that DSF signaling contributes to the virulence of *S. maltophilia*. The pleiotropic effects of loss by mutation *rpfF* are consistent with previous observations in different strains of *S. maltophilia* and species of *Stentotrophomonas* such as *S. rhizophila*, where *rpfF* mutants have altered biofilm formation, extracellular polysaccharide synthesis and virulence (Alvai et al., 2013; Huedo et al., 2015).

The phenotypic effects of *rpfF* mutation in *S. maltophilia* could be reversed by addition of exogenous DSF. Addition of synthetic DSF from *X. campestris* at 1 mM or extracts from wild-type *S. maltophilia* to cultures of the *S. maltophilia rpfF* mutant of an equivalent volume restored wild-type planktonic growth in L medium and restored swimming motility towards wild-type levels (Fig. 4).

**Fig 4.**
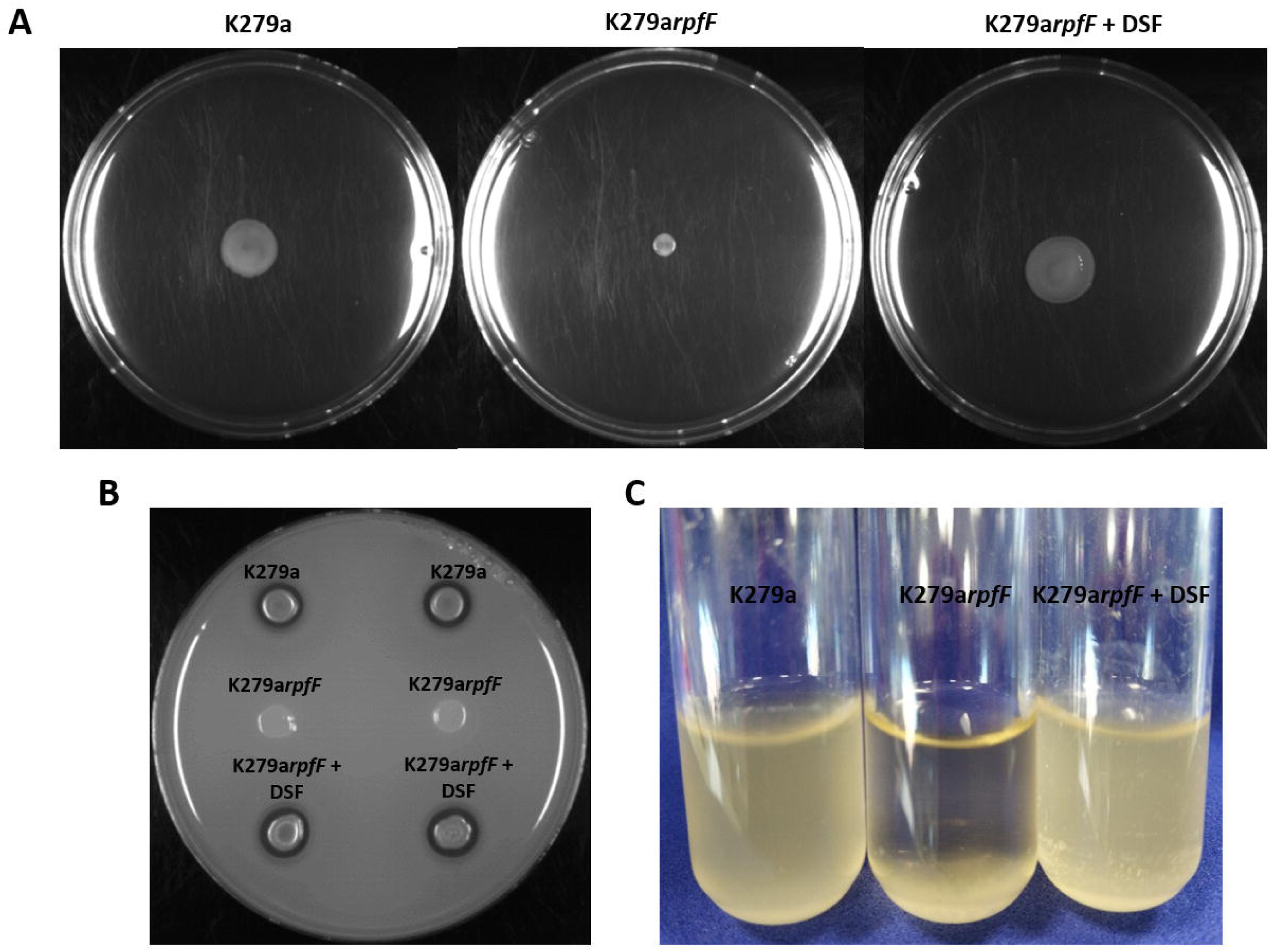
Addition of DSF to agar plates restores swimming motility to the *S. maltophilia rpfF* mutant. The addition of DSF to agar plates restores (A) swimming motility; (B) production of extracellular protease; (C) wild-type planktonic growth in L medium to the *S. maltophilia rpfF* mutant.

## Conclusions

The work in this paper suggests that DSF signaling in *S. maltophilia* has a role in the regulation of a number of functions that contribute to aggregation or biofilm formation and to the virulence of this organism. Our findings thus add to a body of work that indicates a role for cell-cell signaling in the virulence of diverse bacterial pathogens. Furthermore, the work is consistent with recent studies that identify the importance of RpfF proteins and DSF signaling in regulation in other environmental strains of *S. maltophilia* (Alvai et al., 2013; Huedo et al., 2015). Interference with such signaling processes affords a rational approach to aid the treatment of bacterial infections (Ryan et al., 2015). However, one limitation of such an approach is that differences in the operation of Rpf-DSF mediated cell-cell signaling have been reported in different strains of *S. maltophilia* to include both clinical and environmental isolates (Alvai et al., 2013). In this context, a detailed study of DSF signaling and its role in a wider number of *S. maltophilia* isolates is very much warranted.

## Acknowledgements

We thank Robert Ryan, Max Dow and Delphine Caly for initial data, helpful discussions and critical reading of the manuscript.

